# Statistical Quality Scale 6 (SQS-6)

**DOI:** 10.1101/089649

**Authors:** José Fausto de Morais, Gecilmar Pereira Borges, Gilmar Fernandes do Prado

## Abstract

**Context:** The statistical analysis is an important part of the process of assessing the quality of randomized controlled trials, unfortunately it tends to be underestimated or even omitted by editors, in this sense, scales that cover this gap becomes necessary.

**Objective:** To build a definition and a scale for assessing the quality of statistical analysis in randomized controlled trials.

**Methods:** A content analysis on 16 biostatistics texts was considered in the building of the definition and of the scale. The indicators of quality were based on the description and presentation of results of articles.

**Results:** We identify 32 quality indicators grouping in six dimensions: randomization and management of lost; sample size computation; use of mean and median; statistical test; use and interpretation of confidence interval and P-values. The scale range from 0 to 6 and score greater than three identify article with appropriate statistical analysis.

**Conclusions:** The scale presented three dimensions linked to quality of description and three linked to quality of used strategy.

## INTRODUCTION

The randomized controlled trial (RCT) is a research model in which the participants in a study are randomly assigned to an experimental group and control group [1]. The groups are monitored simultaneously and prospectively until we observe the desired outcome. In general the outcome variable is a dichotomy (improve/no improve; survive/die, etc.)

The importance of the RCTs in the evaluating of effectiveness of treatments is known. Many authors [2,3,4] consider the RCTs the most reliable method to evaluate of effectiveness of treatments and one of principal source of information for clinic practice [1,4]. Obviously the report published describing the RCT should have high quality, suggesting higher credibility and acceptance of results by the scientific community [3].

It is important distinguish the difference between quality of trial and quality of its report. Vehagen et al [5] define quality of a trial as “a set of parameters in the design and conduct of a study that reflects the validity of the outcome, related to the external and internal validity and the statistical model used. With regard to the quality of the report, Moher et al [3] defines it as “the provision of information about the design, conduct and analysis of trials.” For the authors, a test well designed and reported, but poorly conducted may receive high score for quality and vice versa.

The first scale to assess quality of a trial/report was published in 1981 by Chalmers *et al* [6]. Until 2001 about 60 scales had been published, unfortunately most of them ignore the third dimension of Verhagen *et al* (the Statistical Model).

Considering that the inappropriate use of statistical analysis may lead to incorrect conclusions and waste of resources [7]; that the quality of report give us an indication of the quality of research [3]; and considering that the first step to building of a scale is the definition of “construct” targeted by the scale [3], this article was prepared with the objective of build a definition and a scale for assessing quality of statistical analysis in RCTs with dichotomous outcome.

## METHODS

In the build of the definition of the statistical quality and of the scale for measure her; we adopted the technique of content analysis [8] on the guidelines contained in a *corpus* with 16 biostatistics texts [7, 9-23]. Briefly, the content analysis was based on the quantification of the occurrence of the variables of interest, allowing classifying them according to the frequency with which they are reported in the texts of the *corpus*. For purpose of this study, we identified and listed the statistical tests or data analysis strategies more frequently used in the texts selected. The analysis was conducted in the following steps:

1. Analyze the texts of the *corpus* (JFM);
2. List the statistics relevant recommendations contained in the *corpus* (JFM);
3. Group recommendations by thematic similarity (JFM);
4. Define labels for grouping (JFM);
5. Build the quality definition from the labels defined (JFM and GPB);
6. Convert each of the groups to a set of quality indicators (JFM);
7. Discuss the proposed indicators to get consensus (JFM and GPB).

## RESULTS

The reading and discussion of texts allowed identifying 32 recommendations applied to statistical analysis of RCTs:

1. Verify if the randomization process includes reference to random numbers table, random number generator or institution that performed the process [9,10].
2. Remember that the results of the research will be questionable if more than 10% of respondents not completing the study [9].
3. In RCTs the analysis by intention to treat should be emphasized [7,10].
4. The text should present the correct name of the statistical test used with a statement that the data meet the conditions for application of the test [11].
5. The comparison groups were established at random and was performed the comparison with baseline [11].
6. Do not use the chi-square test in 2 × 2 tables when identify expected frequency below 5 or if the sample size is less than 20 observations [7,12].
7. Do not make multiple comparisons using the ANOVA procedure without correct the alpha using for example the Bonferroni criteria [13].
8. Do not apply the Student t test for paired data or ANOVA with repeated data in samples with different sizes [14,15,16].
9. Do not use the Student t test or ANOVA for compare variable with ordinal level of measurement [14,15].
10. Do not use the paired Student t test or ANOVA for compare two (or more) independent groups [11,15].
11. Use the Fisher exact test in 2 × 2 tables when some expected frequency is less than 5 or your sample is less than 20 observations [15].
12. You must use the Mann-Whitney U test in the comparison of two groups when comparison variable has ordinal level or it is quantitative and has not Gaussian distribution [15,18].
13. Do not use unusual methods without reference or an explanation of the same [7].
14. If more than one test was used in the study specify which test was applied in each case [7,5]
15. The text should mention the significance level (alpha) and statistical power used in the calculation of “n”. It is usual adopted an alpha of 5% and power of at least 80% [16,17,18,19].
16. In the calculation of “n” to compare two groups using a continuous measure, the text should tell where it came from the estimate of the variability of the measure and must mention the clinically acceptable difference [12,15,17].
17. In the calculation of “n” to compare two groups using prevalence, the text should mention estimative of prevalence in each group [1,7].
18. In the calculation of “n” to estimate of mean or prevalence, the text should mention the confidence level and standard error of the measure of interest [7].
19. Never calculate mean and standard deviation of ordinal level variables, including Likert scales [21].
20. Do not use the standard error (SE) in the place of standard deviation (SD), and do not mention average without reference to standard deviation [11].
21. If variable is ordinal or continuous with non-Gaussian distribution, consider median and quartiles as descriptive measures [11,15].
22. When to use proportion adopts the absolute form, e.g. use 10/20 not 50% [11].
23. Inform the purpose of the study and your primary outcome measure [7,11].
24. Use the exact P value and adopt three decimal places (except when P is less than 0.001. In this case indicate p<0.001) and confidence intervals (in general with 95% confidence) [1,7].
25. Many numbers do not need to be reported with full precision. For instance, use “about 30000” and not 29 942 in the text [11].
26. When categorizing continuous variables explain to reader how the boundaries (or “cut points”) of the categories were determined.
27. Use tables and figures to communicate information, not simply to “store” data. An illustration should help the reader understand the phenomenon [11,15].
28. Remember that a large P-value means that a biggest sample is required to identify difference or effect [11,23].
29. In the 95%CI, the percentage 95% does not indicate that the parameter oscillates between the interval limits with 95% of probability, but that 95% of intervals, with the same sampling error, contain the parameter [22].
30. Remember, small difference between large groups can be statistically significant but clinically meaningless, and large difference between small groups can be clinically important but not statistically significant [11,23].
31. Do not confuse the ‘unit of observation”. For example, in a study with 50 eyes the unit of observation is “eye” and not patient [11].
32. Do not interpret results not statistically significant and low statistical power (n small) as ‘negative’; they are inconclusive [11].

The recommendations produced six groups or conceptual dimensions (D1, D2, D3, D4, D5, and D6) whose labels were named “process of randomization and managing of losses”; “statistical tests”; “calculation of sample size”; “descriptive measures”; “presentation of results”; and “interpretation of results”, respectively. The suggested quality indicators in each dimension are show below:

### D1-Process of randomization and managing of losses

This dimension is composed of items 1, 2 and 3, which can be synthesized in the conditions: “the article should present losses of less than 10%; should adopt the intention-to-treat principle, and should mention the use of the table of random number, random number generator or randomization center”.

### D2-Statistical tests

This dimension is composed by items from 4 to 13 that can be synthesized in the condition: “the article should present the justification (why and for what) for the choice of each statistical test”.

### D3 - The calculation of sample size

This dimension is composed by items from 15 to 18 that can be synthesized in the conditions: “the text should mention the statistical power, the level of significance and the clinically acceptable difference”.

### D4 – The descriptive measures used

This dimension is composed by items from 19 to 22 that can be synthesized in the condition: “the text should use the Median to describe ordinal variable. If the variable is quantitative we can use Mean with Standard Deviation or Median”.

### D5 – The presentation of results

This dimension is composed by items from 23 to 27 that can be synthesized in the conditions: “in the abstract the article should display 95% confidence intervals in conjunction with the p-values, these values must be presented with your exact value except when it is less than 0.001, in which case we should report p<0.001”.

### D6 - Interpretation of results

This dimension is composed by items from 28 to 32 that can be synthesized in the condition: “confidence intervals should not be interpreted in terms of probability (or chance) and neither results with p<0.05 as absence of statistically significant difference”.

Using the aggregate concepts process, described in Cerreto [24], we define: ***Quality of Statistical Analysis is the adequate application of statistics guidelines in: randomization process and management of losses; in determining of sample size; in*** use of descriptive measure; in use of statistical tests; in presentation of results and interpretation of confidence intervals and p-values.

The Table 1 displays the items and scores for scale suggested by analysis, hereinafter referred to as Statistical Quality Scale 6 (SQS6).

**Table 1.** Items and scores for the SQS-6

| Item | Text | Score |
| --- | --- | --- |
| 1 | Was the randomization process and management of losses adequate? | 0=No 1=Yes |
| 2 | Was the description of calculating “n” adequate? | 0=No 1=Yes |
| 3 | Were mean and median used adequately? | 0=No 1=Yes |
| 4 | Does the text explain to the reader why each statistical test was adopted? | 0=No 1=Yes |
| 5 | Were 95%Confidence Intervals used in conjunction with P-values in presentation of results? | 0=No 1=Yes |
| 6 | Were the 95%Confidence Intervals and P-values correctly interpreted? | 0=No 1=Yes |

The items of scale obey the criteria:

<u>Item1</u> - it will receive score 1 if the text reports losses of less than 10%, and clarify whether the process of randomization used a random numbers table, random number generator our central of randomization. It will receive score 0 otherwise.

<u>Item2</u> - It will receive score 1 if text cite statistical power, level of significance and difference clinically acceptable in your description of calculation of “n”. It will receive score 0 otherwise.

<u>Item3</u> - It will receive score 1 if the text display median for ordinal variables or Likert-type scales. When appropriate the text use mean with standard deviation. It will receive score 0 otherwise.

<u>Item 4</u> it will receive score 1 if text explains to reader why each statistical test was selected for study. It will receive score 0 otherwise.

<u>Item 5</u> it will receive score 1 if text displays confidence intervals in conjunction with P-values. These P-values should be presented with their exact value except when it is less than 0.001, in which case we report P <0.001. It will receive score 0 otherwise.

<u>Item 6</u> It will receive score 1 if the confidence intervals are not interpreted in terms of probability (or chance) and no results with P>0.05 are interpreted as no statistically significant difference. It will receive score 0 otherwise.

## DISCUSSION

This study identified six operational guidelines related to the quality of statistical analysis and its description. Three of them associated with the formal description and three associated with the conceptual consistency that supported the strategy of analysis chosen by the author. This is the first study to produce a scale from the perspective of the statistical analysis and its description.

The operational guidelines established for items 1, 2, and 4 of scale are entirely based on the reports and are therefore vulnerable to technical knowledge of the authors, so it does not take into account if the statistical test used was the most appropriate or whether the formula (when used) for calculating of “n” was the most appropriate. Moreover, the guidelines established for items 3, 5, and 6 allow to check failure in use of mean or median and failure in the presentation of results and interpretation of confidence intervals or p-value, regardless of these values are correct or not.

It is important to note that the scale has three items are highly dependent on the reporting of the author (descriptive dimension of the scale) and three items that project the authors’ knowledge of some important statistical concepts (analytical dimension of the scale), in that sense, if we consider as acceptable studies with a score more than three will be ensuring that the article meets, at least, one of the items that measure the statistical knowledge.

Despite the enormous time and energy required to develop a scale [3], we observed that the majority of scales to measure quality of articles, developed until 2001 overlook the statistical analysis dimension. In addition, all items selected for use in these scales are based on acceptance criteria for clinical texts patterns. Although these criteria might be helpful, some of them are based on beliefs of authors, while others are based on empirical evidence [3]. The scale proposed in this study is characterized by the fact that the criteria used in assigning the scores are based on theoretical assumptions in the long time adopted in statistics.

The SQS-6 scale has the virtue of measure some level of statistical knowledge on the part of authors and the defect of being heavily dependent on reporting contained in the articles. Ideally, the raw data were available to the reviewers for the statistical calculations could be revised [25-27], however this proposal is not realistic; since the reviewers are not paid and do not necessarily have the experience required for that service [26,27].

Most of the criteria for evaluation of each dimension of the SQS-6 are abstraction that, in practice, it cannot be verified with the usual mathematical rigor, however, we can use the quality of description of analysis as an estimate of the quality of analysis [3]. This procedure increased the accuracy [28] of the SQS-6 and reinforced its position as tool to assess the statistical quality of RCTs.

The aim of this study was to propose a definition and scale to measure the quality of statistical analysis of RCTs, being outside of his the scope to discuss the psychometric properties (reliability and validity) of the scale. Validation studies in the style of Masuko et al [29], Ferreira et al [30], Oliveira Jr and Morais [31] may be conducted on SQS-6, so it can be used for estimating the statistical quality of comparative studies, especially randomized controlled trials.

## Author Contributions

Study concept and design: JFM and GFP

Acquisition of data: JFM and GPB

Analysis and interpretation of data: JFM and GPB

Drafting of the manuscript: JFM, GFP and GFB

Critical revision of the manuscript: JFM and GFP

Study supervision: GFP Statistical expertise: JFM

All authors read and approved the final manuscript and declare that have no competing interests.

